# Reduction in mRNA expression of the neutrophil chemoattract factor CXCL1 in *Pseudomonas aeruginosa* treated Barth Syndrome B lymphoblasts

**DOI:** 10.1101/2023.04.18.537385

**Authors:** Hana M. Zegallai, Kangmin Duan, Grant M. Hatch

**Affiliations:** Children’s Hospital Research Institute of Manitoba, Department of Pharmacology & Therapeutics; Department of Oral Biology, University of Manitoba, Winnipeg, Manitoba, Canada, R3E0T6

**Keywords:** Barth Syndrome, X-linked genetic disease, human B lymphoblasts, *TAFAZZIN*, cardiolipin, chemokine (C-X-C motif) ligand 1, CXCL1, *Pseudomonas aeruginosa*, bacterial activation, immune biology

## Abstract

Barth Syndrome (BTHS) is a rare X-linked genetic disease caused by a mutation in *TAFAZZIN*, a cardiolipin transacylase. Approximately 70% of patients with BTHS exhibit severe infections due to neutropenia. However, neutrophils from BTHS patients have been shown to exhibit normal phagocytosis and killing activity. B lymphocytes play a crucial role in the regulation of the immune system and when activated secret cytokines known to attract neutrophils to sites of infection. We examined expression of chemokine (C-X-C motif) ligand 1 (CXCL1), a known chemotactic for neutrophils, in Epstein-Barr virus transformed control and BTHS B lymphoblasts. Age-matched control and BTHS B lymphoblasts were incubated with *Pseudomonas aeruginosa* for 24 h and then cell viability, CD27+, CD24+, CD38+, CD138+ and PD1+ surface marker expression and CXCL1 mRNA expression determined. Cell viability was maintained in lymphoblasts incubated with a ratio of 50:1 bacteria:B cells. Surface marker expression was unaltered between control and BTHS B lymphoblasts. In contrast, CXCL1 mRNA expression was reduced approximately 90% (*p*<0.05) in untreated BTHS B lymphoblasts compared to control cells and approximately 70% (*p*<0.05) in bacterial treated BTHS B lymphoblasts compared to control cells. Thus, naïve and bacterial-activated BTHS B lymphoblasts exhibit reduced mRNA expression of the neutrophil chemoattractant factor CXCL1. We suggest that impaired bacterial activation of B cells in some BTHS patients could promote immune dysfunction, and this may contribute to infections.

## Introduction

Barth Syndrome (BTHS) is a rare X-linked genetic disease of young boys [1-7]. It is caused by mutation in the gene *TAFAZZIN*. The protein product tafazzin is a transacylase enzyme responsible for remodeling of cardiolipin [8]. Cardiolipin is a key mitochondrial phospholipid that is essential for activation of many enzymes of the electron transport chain. As a result, cells from BTHS patients have reduced ability to produce ATP from oxidative phosphorylation [1-7]. This leads to development of cardiomyopathy, skeletal myopathy, growth retardation, 3-methylglucaconic aciduria, and neutropenia.

Although cardiomyopathy is the main pathological problem in BTHS, as most boys develop heart failure, many of these boys (approximately 70%) exhibit severe infection due to neutropenia [9]. Interestingly, neutrophils isolated from BTHS patients have been shown to exhibit normal phagocytosis and killing activity [10]. The neutropenia of BTHS is typically treated with granulocyte-colony stimulating factor to restore neutrophil numbers but in some cases infections persist [9]. B lymphocytes are known to influence and modulate neutrophil function through synthesis and secretion of products such as cytokines and chemokines. For example, B lymphocytes produce chemokine (C-X-C motif) ligand 1 (CXCL1) which is a known chemotactic for neutrophils [11]. In the current study, we examined if BTHS B lymphoblasts exhibited impaired ability to express the neutrophil chemoattractant factor CXCL1. We show that both naïve and *Pseudomonas aeruginosa*-stimulated BTHS B lymphoblasts exhibit impaired mRNA expression of CXCL1. We hypothesize that impaired bacterial activation of B lymphocytes could promote immune dysfunction in some BTHS patients and this may contribute to infections.

## Materials and Methods

### Materials

All work was performed with approval from the University of Manitoba Environmental Health and Safety Office (Biological Safety Project Approval Certificate #BB0044-2). Epstein-Barr virus transformed human control lymphoblasts and Epstein-Barr virus transformed BTHS patient lymphoblasts were obtained from the Coriell Institute for Medical Research (Camden, NJ, USA). The cells that were used in the study are outlined in **Table 1**. RPMI 1640 media, Fetal bovine serum (FBS), antibiotic-antimycotic, and propidium iodide were obtained from Life Technologies Inc. (Burlington, ON, Canada). Anti-CD19-APC, anti-CD24-Pacific blue, anti-CD27-perCP, anti-CD38-APC-CY7, anti-CD138-PE, and anti-PD1-APC antibodies for flow cytometry analysis were purchased from BD Biosciences (Seattle, WA, USA). Unless otherwise indicated, all other reagents used were of analytical grade and were obtained from either ThermoFisher Scientific (Winnipeg, MB) or Sigma-Aldrich (Oakville, ON).

**Table 1.**
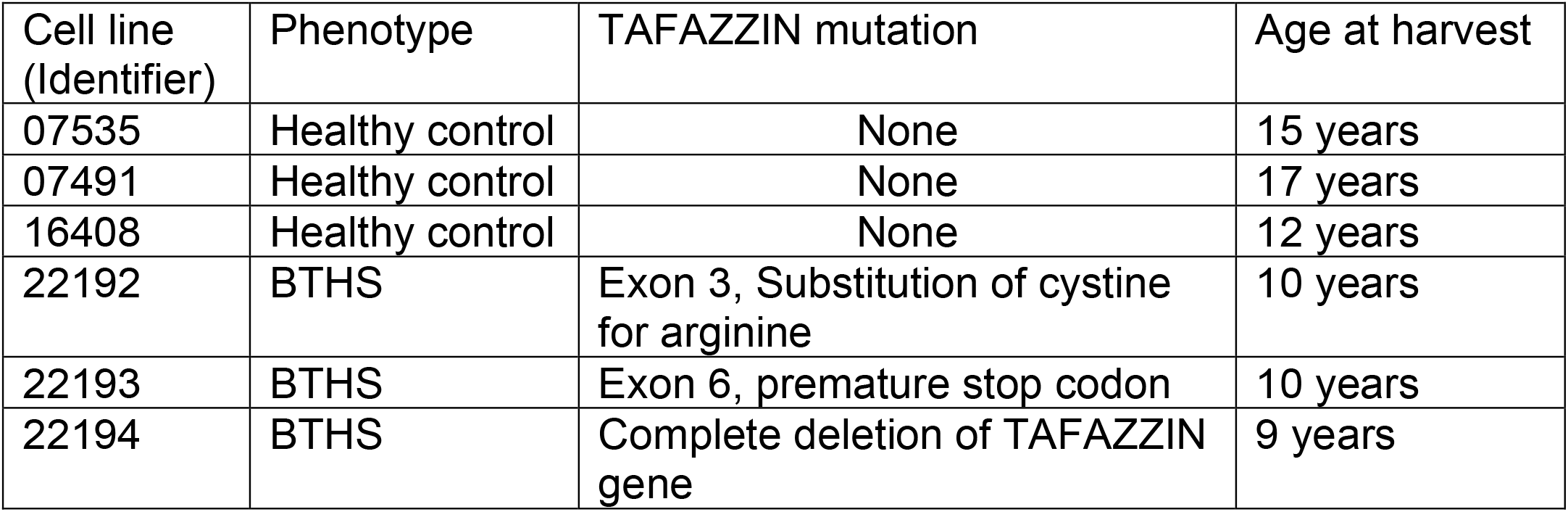
Patient cell lines used in this study.

### Cell culture and bacterial stimulation

All work was performed with approval from the University of Manitoba Environmental Health and Safety Office (Biological Safety Project Approval Certificate #BB0044-2). Cells were grown in RPMI-1640 medium supplemented with 15% FBS and 1% antibiotic-antimycotic at 5% CO_2_ at 37°C in a Thermo Scientific CO_2_ incubator HEPA Class 200. The medium was replaced every 48 h and the cells were passaged every five days. Cells were pelleted by centrifugation at 1400 rpm for 10 mins at room temperature. Cells were then washed twice with phosphate buffered saline (PBS) prior to experimental stimulation. Control and BTHS lymphoblasts were incubated plus or minus live *Pseudomonas aeruginosa* bacteria (at exponential phase) in a 50:1 ratio of bacteria:human cells for 24 h. After treatment, the cells were harvested by centrifugation as above and washed twice with PBS prior to further analysis.

### Relative Gene Expression

RNA was isolated from control and BTHS patient lymphoblasts using RNeasy Mini Kit and Qiashredder homogenizer columns. RT-PCR was performed using the QuantiTect® Probe RT PCR Kit and the double stranded DNA stain SYBR green as indicated by the manufacturer. Relative gene expression analysis was calculated using the 2-ΔΔCt method. The primers used for RT-PCR detection were as follows: CXCL1: forward, 5′-GAACATCCAAAGTGTGAACGTGAAG-3′; reverse, 5′-TTCAGGAACAGCCACCAGTGAG-3′; β-actin: forward, 5′-GG CGGCACCACCATGTACCCT-3′; reverse, 5′-AGGGGCCGGACTCGTCA TACT-3′. All primers were obtained from Integrated DNA Technologies (Coralville, IA).

### Cell viability and surface marker expression analysis

Briefly, after 24 h of treatment with live bacteria, the lymphoblasts were centrifuged at 1400 rpm for 10 min. The pellets were washed with PBS and suspended in 100 μl of PBS. Untreated and bacterial treated lymphoblasts were stained with propidium iodide (5 μg/ml) for 5 min in the dark at 4°C. This was followed by flow cytometry analysis (see below). Surface marker expression was measured in untreated and bacterial treated lymphoblasts by staining the cell surface with anti-CD19-APC, anti-CD24-Pacific blue, anti-CD27-perCP, anti-CD38-APC-CY7, anti-CD138-PE, and anti-PD1-APC as per the manufacturer’s instructions. Cells were then analyzed by flow cytometry at the Flow Cytometry Core Facility in the Rady Faculty of Health Sciences, University of Manitoba, using a BD FACS Canto II instrument. FlowJo software was used for data analysis.

### Statistical analysis

All experimental results were expressed as mean ± SD. A two tailed unpaired Student’s t test was performed to compare between groups. The level of significance was defined as p<0.05.

## Results

Initially we examined cell viability of control and BTHS lymphoblasts treated with *Pseudomonas aeruginosa*. Control and BTHS lymphoblasts were incubated for 24 h in the absence or presence of a 50:1 ratio of bacteria:B cells and then stained with propidium iodide and cell viability determined using flow cytometry analysis. Cell viability was maintained in both control and BTHS B lymphoblasts incubated for 24 h with *Pseudomonas aeruginosa* (**Fig. 1**).

**Figure 1.**
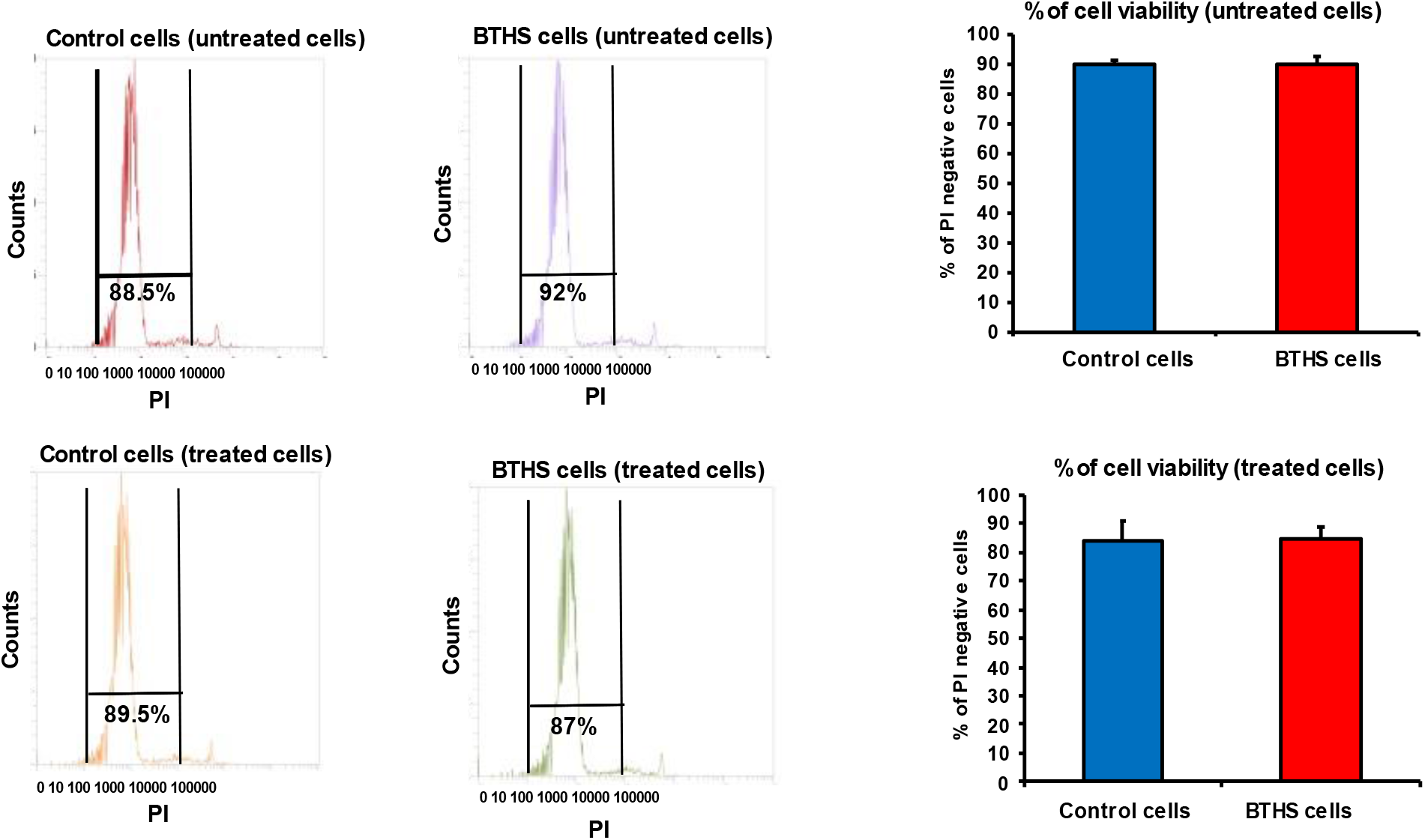
Cell viability of control and BTHS lymphoblasts incubated with bacteria. Control and BTHS lymphoblasts were incubated for 24 h plus or minus *Pseudomonas aeruginosa* and cell viability determined using flow cytometry analysis as described in Materials and Methods. Representative flow cytometry plots are on the left. Data represent the mean ± SD, n=3.

Activation of B lymphocytes with bacteria results in the expression of surface markers such as CD138+. We next examined expression of surface markers on control and BTHS lymphoblasts treated with *Pseudomonas aeruginosa*. Control and BTHS lymphoblasts were incubated for 24 h with *Pseudomonas aeruginosa* and surface expression of CD27+, CD24+, CD38+, CD138+ and PD1+ determined using flow cytometry analysis. Surface expression of CD27+, CD24+, CD38+ and PD1+ was unaltered between untreated control and BTHS lymphoblasts (**Fig. 2, Suppl. Fig. 1**). In addition, when cells we treated with *Pseudomonas aeruginosa* surface expression of CD27+, CD24+, CD38+ and PD1+ was unaltered between control and BTHS lymphoblasts. As expected CD138+ was minimally expressed in unstimulated lymphoblasts. When cells were treated with *Pseudomonas aeruginosa* surface expression of CD138+ was upregulated in both control and BTHS lymphoblasts.

**Figure 2.**
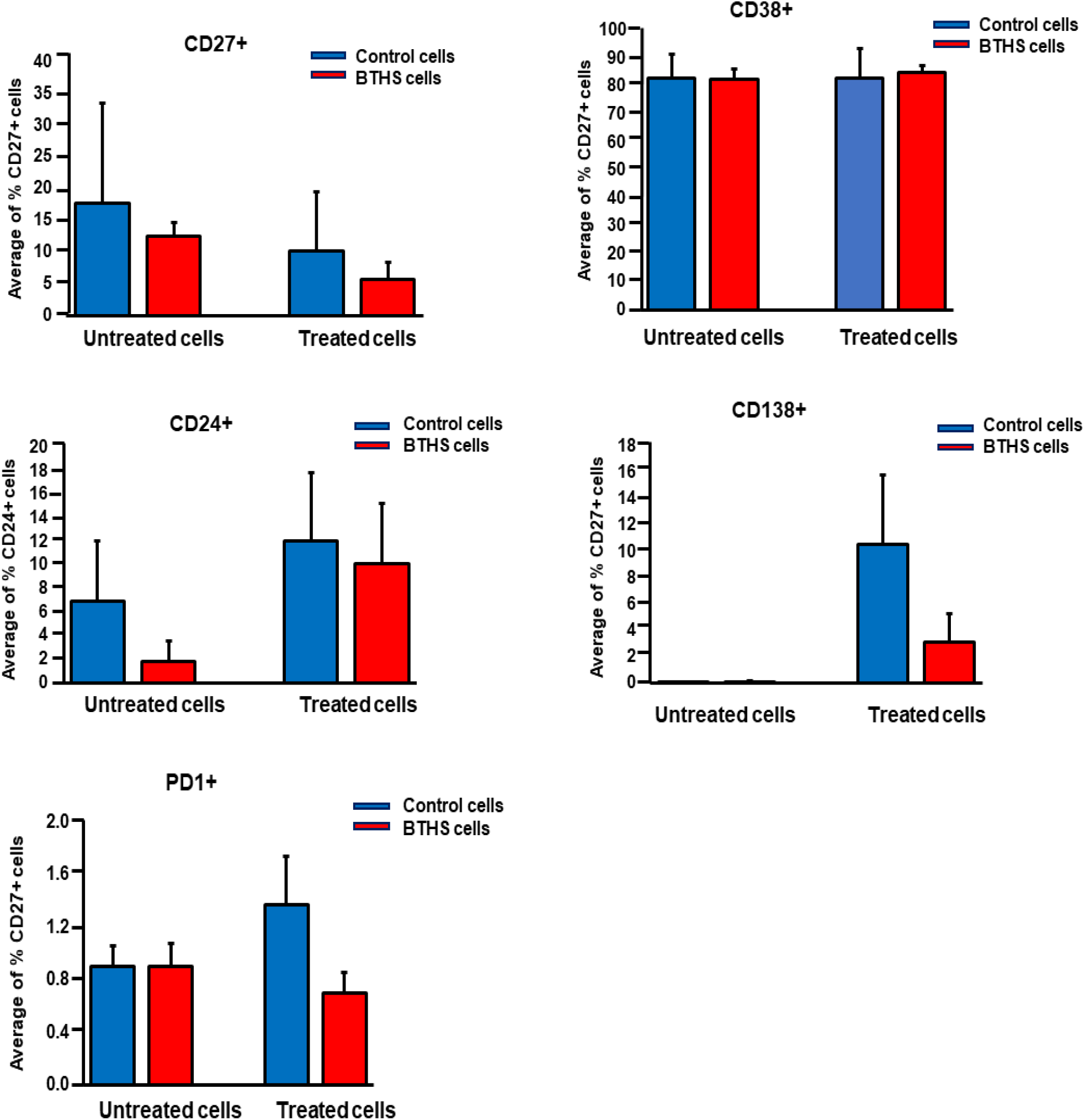
Surface marker expression in control and BTHS lymphoblasts incubated with bacteria. Control and BTHS lymphoblasts were incubated for 24 h plus or minus *Pseudomonas aeruginosa* and surface marker expression of CD27+, CD24+, CD38+, CD138+ and PD1+ determined using flow cytometry analysis as described in Materials and Methods. Data represent the mean ± SD, n=3.

Next, we examined mRNA expression of CXCL1 in control and BTHS lymphoblasts treated with *Pseudomonas aeruginosa*. The relative mRNA expression of CXCL1 in untreated naïve BTHS lymphoblasts was 90% lower compared to untreated naïve control lymphoblasts (**Fig. 3A**). In addition, the relative mRNA expression of CXCL1 in *Pseudomonas aeruginosa* treated BTHS lymphoblasts was 70% lower compared to *Pseudomonas aeruginosa* treated control lymphoblasts (**Fig. 3B**). When control cells were compared, relative CXCL1 mRNA expression was elevated 7-fold by *Pseudomonas aeruginosa* treatment compared to untreated control cells (**Suppl. Fig. 2A**). In contrast, when BTHS cells were compared, relative CXCL1 mRNA expression was elevated by only 40% in *Pseudomonas aeruginosa* treated BTHS cells compared to untreated BTHS cells (**Suppl. Fig. 2B**) indicating a reduced ability of BTHS B cells to express CXCL1 mRNA compared to control cells. Thus, naïve and *Pseudomonas aeruginosa* stimulated BTHS B lymphoblasts exhibit impaired mRNA expression of CXCL1.

**Figure 3.**
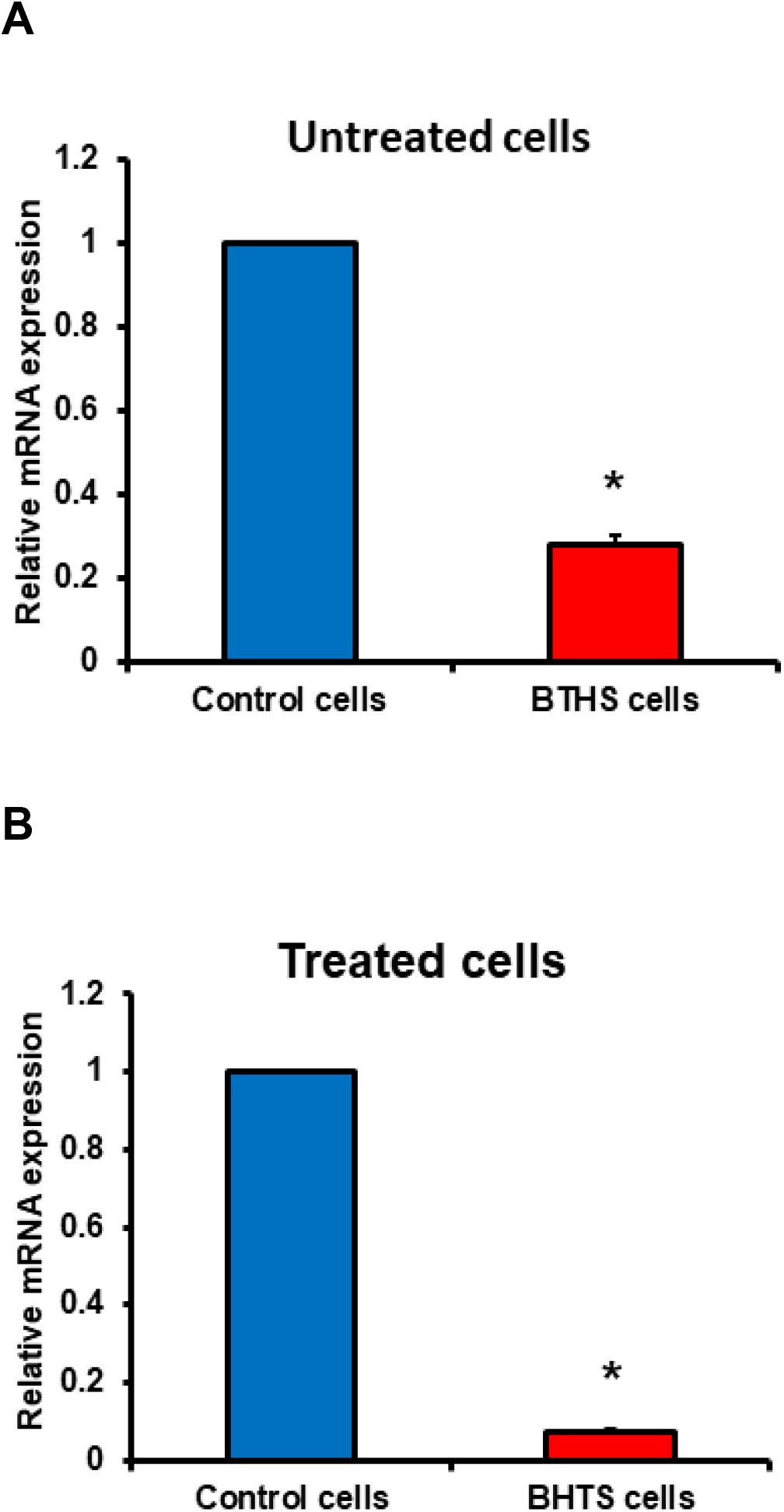
CXCL1 expression is reduced in bacterial treated BTHS patient lymphoblasts. Control and BTHS lymphoblasts were incubated for 24 h plus (treated) or minus (untreated) *Pseudomonas aeruginosa* and CXCL1 mRNA expression determined as described in Materials and Methods. Data represent the mean ± SD, n=3. *p<0.05.

## Discussion

Tafazzin deficiency is known to negatively impact the function of a broad range of immune cells including mesenchymal stem cells, mast cells, CD8+ T cells and B cells [12-15]. Crosstalk between neutrophils and B cells occurs in several diseases including rheumatoid arthritis and antineutrophil cytoplasmic antibody-associated vasculitis [16,17]. CXCL1 is a CXC family member that acts as a key chemoattractant for immune cells, especially neutrophils and other non-hematopoietic cells to sites of injury or infection and plays a key role in regulation of immune and inflammatory responses [18,19]. A previous study had indicated that neutrophils isolated from BTHS patients exhibited normal phagocytosis and killing activity [10]. Moreover, a study examining the clinical data, routine blood counts, and responses to granulocyte-colony stimulating factor therapy from 88 affected BTHS boys concluded that susceptibility to infections was due only in part to neutropenia since in some instances infection occurred despite consistent prevention of neutropenia by granulocyte-colony stimulating factor therapy [9]. A more recent study examined myeloid progenitor development within the fetal liver of TAFAZZIN knockout animals as well as within the adult bone marrow of wildtype recipients transplanted with TAFAZZIN knockout hematopoietic stem cells [20]. Although TAFAZZIN knockout neutrophils demonstrated the expected differences in cardiolipin maturation no significant differences in neutrophil development or neutrophil function, including the production of cytokines, was observed.

In the current study, we observed mRNA expression of CXCL1 in both naïve and *Pseudomonas aeruginosa*-activated control and BTHS lymphoblasts. However, the degree of CXCL1 mRNA expression was significantly lower in naïve or bacterial-activated BTHS lymphoblasts compared to naïve or bacterial-activated control lymphoblasts indicating an impaired ability of BTHS patient B lymphoblasts to express CXCL1.

The presence of CD138+ expression on B lymphocytes is indicative of an active antibody secreting phenotype of B cells [21]. Interestingly, although *Pseudomonas aeruginosa* treatment resulted in expression of CD138+ in both control and BTHS cells, the degree of expression appeared to be lower in BTHS cells than control cells, albeit this was not statistically significant. These results are consistent with the recent identification of a BTHS patient with persistent B cell lymphopaenia and hypogammaglobulinaemia [22]. The patient experienced repeated bacterial and viral infections and required subcutaneous immunoglobulin replacement for ongoing hypogammaglobulinaemia. Thus, it is possible that reduced CD138+ expression on bacterial activated B lymphocytes might be linked to attenuated antibody production in some BTHS patients.

A limitation of our study is the use of Epstein-Barr virus transformed control and BTHS lymphoblasts as opposed to primary human BTHS B lymphocytes. Epstein-Barr virus transformation of primary B lymphocytes results in differential expression of several genes compared to that of normal human B lymphocytes [11]. These include 22 up-regulated genes and 16 down-regulated genes (one of which is CXCL1) which control processes such as the cell cycle, mitosis, the cytokine-cytokine pathway and the immunity response thus hindering host immune function and secretion of cytokines. We recently observed that Epstein-Barr virus transformation of control and BTHS human B lymphocytes exhibited impaired responsiveness to lipopolysaccharide-mediated or CpG-DNA-mediated surface marker expression [23]. Thus, the responsiveness of the patient control and BTHS Epstein-Barr virus transformed lymphoblasts to *Pseudomonas aeruginosa* bacterial activation may be already impaired in comparison to non-transformed naïve patient B lymphocytes.

In summary, we have observed reduced CXCL1 mRNA expression, a known chemoattractant of neutrophils, in naïve and in *Pseudomonas aeruginosa*-activated B lymphoblasts from BTHS patients. We hypothesize that a reduced expression of CXCL1 by B lymphocytes may, in part, help to explain why BTHS boys suffer from infections even when neutrophil counts are at or near normal levels.

## Acknowledgements

We thank Marilyne Vandel for technical assistance and Dr. Christine Zhang for flow cytometry analysis. H.M.Z. was supported by a Libyan North American Scholarship. This work was supported by grants from the Natural Sciences and Engineering Research Council of Canada (NSERC) to G.M.H (RGPIN-05368-2019) and to K.D. (RGPIN-05864-2019).

---

**Supplementary Figure 1.**
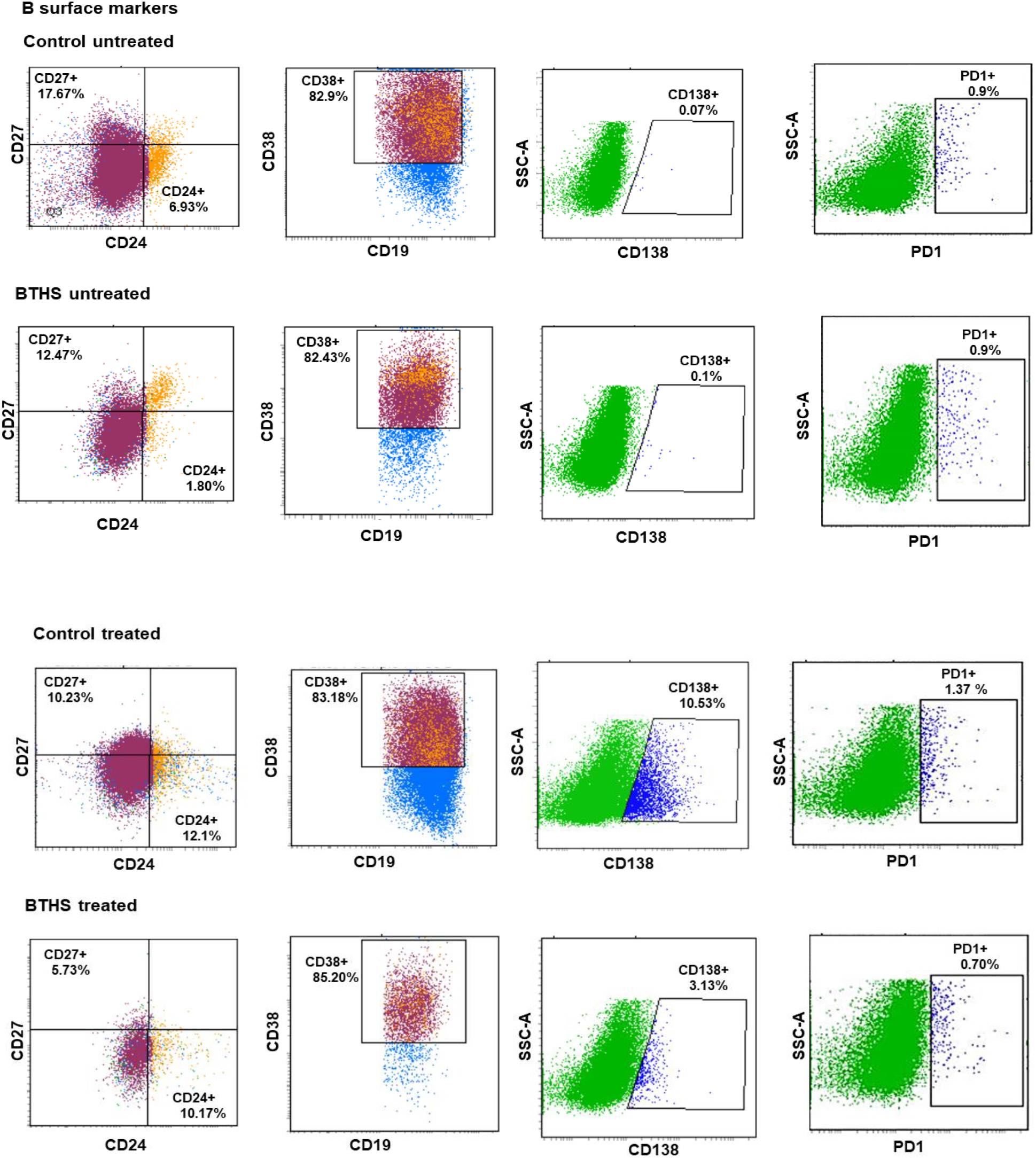
Representative flow cytometry images of surface markers expressed in untreated and *Pseudomonas aeruginosa* treated control and BTHS lymphoblasts.

**Supplementary Figure 2.**
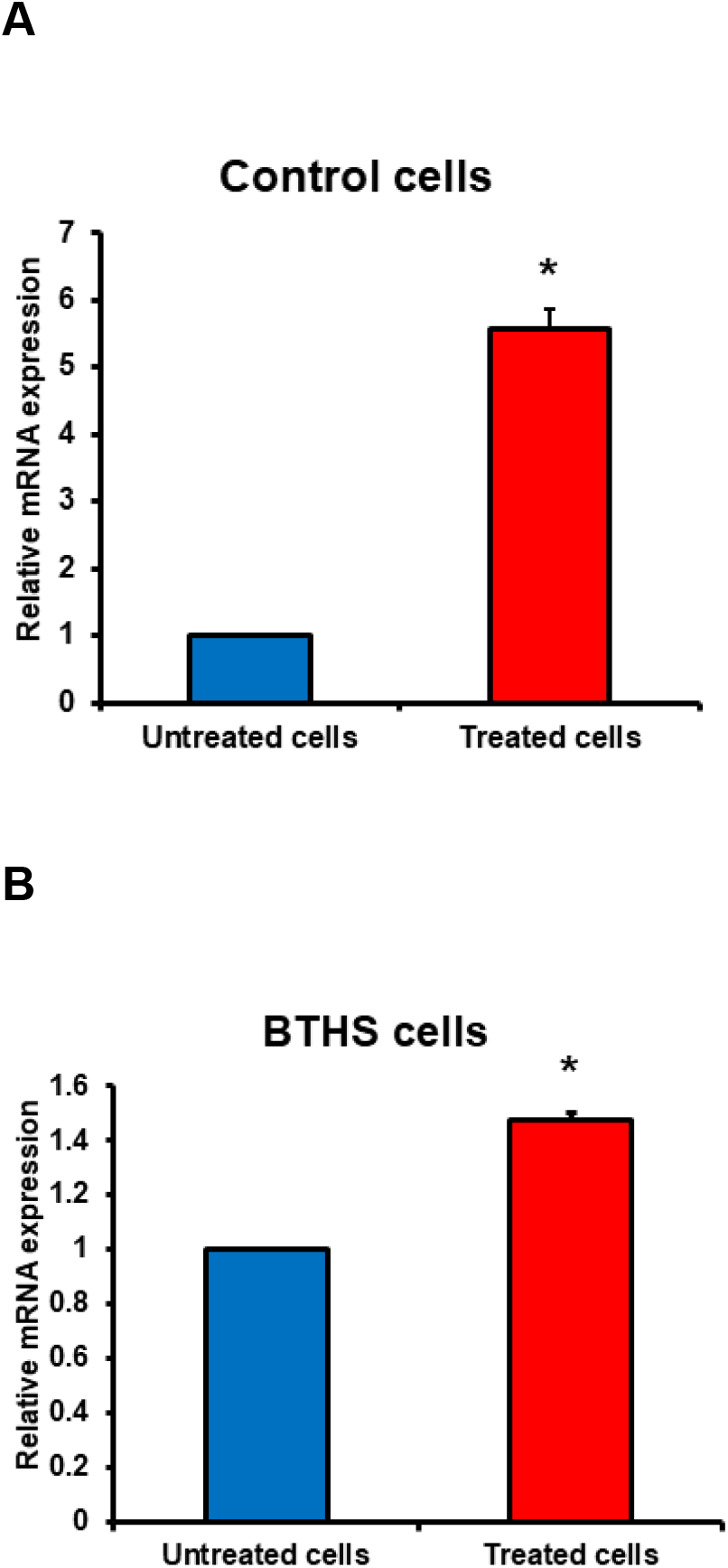
Relative CXCL1 mRNA expression in *Pseudomonas aeruginosa* treated control cells compared to untreated control cells (**A**) and in *Pseudomonas aeruginosa* treated BTHS cells compared to untreated BTHS cells (**B**). n=3, *p<0.05.

